# Coevolution-driven method for efficiently simulating conformational changes in proteins reveals molecular details of ligand effects in the β2AR receptor

**DOI:** 10.1101/2023.07.20.549854

**Authors:** Darko Mitrovic, Yue Chen, Antoni Marciniak, Lucie Delemotte

## Abstract

With the advent of AI-powered structure prediction, the scientific community is inching ever closer to solving protein folding. An unresolved enigma, however, is to accurately, reliably and deterministically predict alternative conformational states that are crucial for the function of e.g. transporters, receptors or ion channels where conformational cycling is innately coupled to protein function. Accurately discovering and exploring all conformational states of membrane proteins has been challenging due to the need to retain atomistic detail while enhancing the sampling along interesting degrees of freedom. The challenges include but are not limited to finding which degrees of freedom are relevant, how to accelerate the sampling along them, and then quantifying the populations of each micro- and macrostate. In this work, we present a methodology that finds the relevant degrees of freedom by combining evolution and physics through machine learning and apply it to the β2 adrenergic receptor conformational sampling. In addition to predicting new conformations that are beyond the training set, we have computed free energy surfaces associated with the protein’s conformational landscape. We then show that the methodology is able to quantitatively predict the effect of an array of ligands on the β2 adrenergic receptor activation, and that the full conformational landscape, including states related to biased signaling, is discovered using this procedure. Lastly, we also stake out the structural determinants of activation and inactivation pathway signaling through different ligands.

## Introduction

From receptors to transporters, cycling between conformational states is how many membrane proteins conduct their function. The energetic landscapes of these proteins are often also modulated by the binding of different ligands as a response to external stimuli, creating a complex mosaic of different behaviors in different conditions. ^1–4^ As fascinating a target this makes them for scientific study, it is often difficult to trap a protein in a certain conformation for long enough to study the structural and functional characteristics, let alone solve a structure in all available conformational states.^5–7^ This leaves many potentially functionally relevant conformational states undiscovered, and insights about functional experiments may be misguided. In addition, with the advent of the structure prediction revolution recently spearheaded by machine-learning models such as Alphafold2 and trRosetta, it has become easier than ever to predict structures of proteins that are difficult to solve experimentally. ^8–10^ Though successful in predicting folds, the question of which conformational state is predicted is not clear. There is an inherent problem with the lack of balanced datasets for different conformational states in many families, which means that models would be discouraged to predict potentially correct alternative states in favor of the ones that are being trained on.^11, 12^ In addition, transition states and the kinetically accessible paths between alternative states are not captured experimentally and are thus impossible to train on thus far.

To probe the short-timescale (<10ns) dynamics of conformational change effectively, one can employ molecular dynamics (MD) simulations, where one parametrizes each atomic interaction with a forcefield, and evaluates Newton’s equations of motion in the time domain to simulate a protein structure. Due to the high computational complexity, large systems over long timescales are difficult to simulate in equilibrium MD, a regime in which many important processes such as receptor activation, membrane transport or ion conduction fall. One approach to prolong the timescales of MD simulations is to enhance the sampling by perturbing the Hamiltonian adaptively based on historical sampling to encourage sampling of new regions and discourage sampling of already visited regions.^13^ In this work, we utilize the accelerated weight histogram (AWH) algorithm,^14^ which adaptively changes a bias potential along a predefined collective variable (CV), which in turn is a function of the atomic coordinates in the system (see methods). Since the added bias of the AWH method is known at all times, it is possible to calculate the convolved free energy surface of a bias-free simulation based on the measured probability distribution. This feature makes it possible to evaluate the energetic stability of the explored conformational space.

Finding an appropriate set of collective variables to sample is crucial for this algorithm to converge and properly sample all functionally relevant conformational states. Learning the CV while having no observation of the motion is difficult, and our previous efforts have been mostly centered on a supervised learning approach.^15, 16^ There are however some general principles for designing a good CV: i) a good CV should describe the conformational change such that all relevant states are separable and uniquely correspond to different points along the CV.^17^ ii) a good CV for AWH or other adaptive biasing methods should contain the degrees of freedom that are responsible for the highest energetic or kinetic barriers. ^13^

An essential hurdle to overcome is that the space of degrees of freedom is very large (3N) and searching all possible configurations would be intangible for very large systems, such as proteins or other biomolecules. Thus, it is clear that we need a way to reduce the space of possible configurations by means other than structural information deposited in the PDB. We previously introduced a supervised coevolution-driven methodology to predict conformational states across entire families; however, this required at least one solved structure in each conformational state.^15^ This type of comparison between structures and coevolutionary information enables the identification of potential so-called false positive coevolving pairs (corresponding to coevolving residues not in contact in a reference structure). We hypothesize that those can represent contacts formed in other conformational states. Since evolutionary information is agnostic of whatever sparse structural information is available and tightly coupled to function, we wondered whether exploring the space of possible contacts from coevolving residue pairs could aid in discovering new states. We reasoned that if we restricted all possible conformations based on kinetic accessibility and thermodynamics as defined by physical force fields in simulations while enhancing the sampling along CVs containing evolutionary information, we could find new, functionally relevant conformational states. The coevolving residue pairs thus provide a convenient dimensionality reduction transformation, whereas the relevant hypervolume to be explored can be determined by an energy-based sampling algorithm, such as MD simulations.

To benchmark this methodology, we chose a family of proteins where it is possible to extract coevolutionary information from the multiple sequence alignments, and where the conformational landscape is well-defined by experimental structures. One of the largest and most pharmaceutically relevant families of membrane proteins are G-protein coupled receptors (GPCRs).^18, 19^ The energetics of these receptors are finely tuned to respond to a multitude of external ligands, from drug-like molecules to lipids or peptides. Thanks to their importance in drug development and signaling pathways, there are a plethora of structures available in various states, making them the perfect candidate to validate our methodology on. We chose the β2-adrenergic receptor (β2AR) as our focus because of the abundance of both structural and functional information in the form of deposited PDB structures and activity experiments. As this system has been a subject of numerous computational studies, we considered it an appropriate target for benchmarking and for comparison of the timescales of convergence. We further validated our methodology by simulating the β2AR bound to nine different ligands with experimentally determined activities and structurally resolved binding modes. The resulting free energy landscapes evaluated that our method is able to sample functional conformational states and accurately predict the efficacies of β2AR ligands, providing insights into the molecular mechanism of activation, inactivation and biased agonism.

## Methods

### Coevolution analysis and false positive detection

Firstly, the multiple sequence alignment (MSA) for the GPCR class A family was downloaded from the PFAM database, now integrated with Interpro, PFAM ID: PF00001, name: 7TM_1, since this was the family to which the target protein belonged. As to not divert the model’s attention from the most conserved regions, we modified the MSA and the training parameters accordingly. Specifically, we filtered positions with more than 20% gaps, and assigned an effective weight to each sequence equal to 1/N_0.9_, where N_0.9_ is the number of sequences with over 90% identity. Then, the MSA was used as a training set for direct coupling analysis (DCA), with the goal to fit a Potts model in the pseudo-maximum likelihood sense (equation 1)

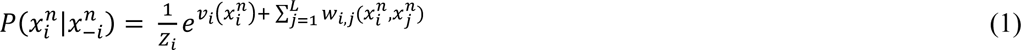

where the conditional probability encodes the position-wise (i, j) information from the entire MSA given the model parameters. The model parameters correspond to column (position)-specific fields 𝑣 and column pair-specific couplings 𝑤. The actual loss function that is optimized is then the natural logarithm of this expression, summed over all sequences n and positions i, and weighted sequence-wise by the 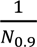 metric. The loss function was optimized using the Adam optimizer for 300 iterations as implemented in TensorFlow, with a learning rate of 0.0001 and parameters initialized randomly by drawing from a normal distribution of mean 0 and standard deviation 1. The code was adapted in the described manner from the original GREMLIN implementation. (See https://github.com/sokrypton/GREMLIN_CPP/blob/master/GREMLIN_TF_v2_BETA.ipynb)

The pairwise coupling parameter w is in the form of a 20x20 matrix, which is then summed over all axes to yield a scalar value of the coevolution strength for a certain (i,j) pair. To reduce the row-wise noise, an average product correction (APC) correction is applied so that comparable scores are attained.

The coevolution scores are then overlaid with the physical contact map of the target protein’s starting structure, and the false positives are identified and clustered based on coevolution strength (see the exploration simulation section for details). We have made this script available on the attached Zenodo project which can be found by following this link https://doi.org/10.5281/zenodo.8164731.

### MD simulations

All Molecular Dynamics (MD) simulations were carried out in GROMACS2021. The simulation systems containing the β2AR in the active state (PDB ID 3P0G) bound to different ligands were prepared using the CHARMM-GUI membrane builder, and the ligands were parametrized using the built-in cgenff procedure available in CHARMM-GUI. Two histidines (H172^4.64^ and H178^4.70^) were protonated at their epsilon positions, and the rest of the histidines were left protonated at the delta position. Furthermore, a Na^+^ ion was placed at the sodium binding site 2.50 and left unprotonated, as done in earlier work.^20^ The systems contained the protein embedded in a POPC bilayer plunged in a 0.1M KCl solution. The initial periodic boundary condition (PBC) box was 85x85x94 Å^3^, ensuring at least 12.5 Å of water molecules between the protein and the PBC box end at least 10 lipid molecules between each PBC copy of the protein. The force field used was CHARMM36m for protein and lipids, and TIP3P for water. The models were equilibrated using the default CHARMM-GUI scheme with one minimization step, and 6 100ps restraint cycles with gradually released restraints in the NPT ensemble, followed by a production simulation of 10ns. The simulations were carried out using a 2-fs time step. The target temperature and pressure were set to 303.15K and 1 bar respectively and maintained by a Nose-Hoover thermostat (coupling separately protein, lipids and solvent) and the recently implemented C-rescale barostat with semi-isotropic coupling (p=5.0, compressibility 4.5*10-5). Hydrogen bonds were constrained using the linear constraint solver (LINCS), and long-range electrostatics were accounted for using the particle mesh Ewald (PME) method beyond the 12Å electrostatic cutoff. A neighbor-list cutoff was used for vdw interactions with r_vdw_=12Å and a switching function starting at 10Å.

### Exploration Simulations

The non-equilibrium exploration simulations for the apo system used the same parameters as described above, but the Hamiltonian of the system was further modified after equilibration. The false positives identified by aligning the coevolution map and the contact map calculated from the structure were clustered into three clusters based on coevolution strength with the K-means algorithm, the minimum-distance atom pairs were used to define individual distances by the pull code in GROMACS. Then, three transformation pull coordinates corresponding to the three clusters (determined by the mean-shift algorithm) expressed the mean of all intra-cluster distances respectively for each of the three clusters. Then, umbrella potentials were applied to each transformation pull coordinate in four independent replicas, in order to push the system towards an alternative state. The force constant used was 200kJ/(mol nm^2^) per contact and the target was 0.2 nm . After 10ns of pulling, the added potential was switched off and an additional 10ns of unrestrained MD simulations was run. The resulting trajectories were then used to train a SVM classifier with a linear kernel to distinguish the ensembles from the equilibrium simulations based on the reference structure 3P0G and end-state of the non-equilibrium pulling simulations. The top 20 contacts were then used to define two CVs through a weighted sum - one with the positive and one with the negative coefficients. Ultimately, these were then used to define the CV space for the AWH simulations. The exploration simulations were not repeated for each ligand system, but rather the CV generated for the apo system were used for all ligand-bound systems, ensuring that a common projection in a set of low-dimensional projections. The necessary files to reproduce these simulations can be found in the associated Zenodo project (see the following link https://doi.org/10.5281/zenodo.8164731).

### AWH simulations

The AWH simulations were run with 4 walkers, two starting from the equilibrated reference structure based on 3P0G and two starting from the end-state as explored by the non-equilibrium pulling simulations. All necessary input files including tpr files are available at the Zenodo project (https://doi.org/10.5281/zenodo.8164731) As the explored state may be far from equilibrium, we set the growth factor parameter to 2.0, which helped prolong the number of initial sweeps required for the simulations to initiate the Wang-Landau algorithm. As mentioned in our previous work, we have experienced that free energy estimates stemming from this modified parameter produce long-term reliability and flatter distributions without walkers getting stuck. Furthermore, while the cover diameter, energy cutoff, number of steps between each AWH sample and number of samples between updates were respectively set to 0.4 nm, 120 kJ/mol, 10 and 10 for all simulations, parameters for individual simulations were fine-tuned to allow for convergence on faster timescales (see Table 1).

**Table 1.**
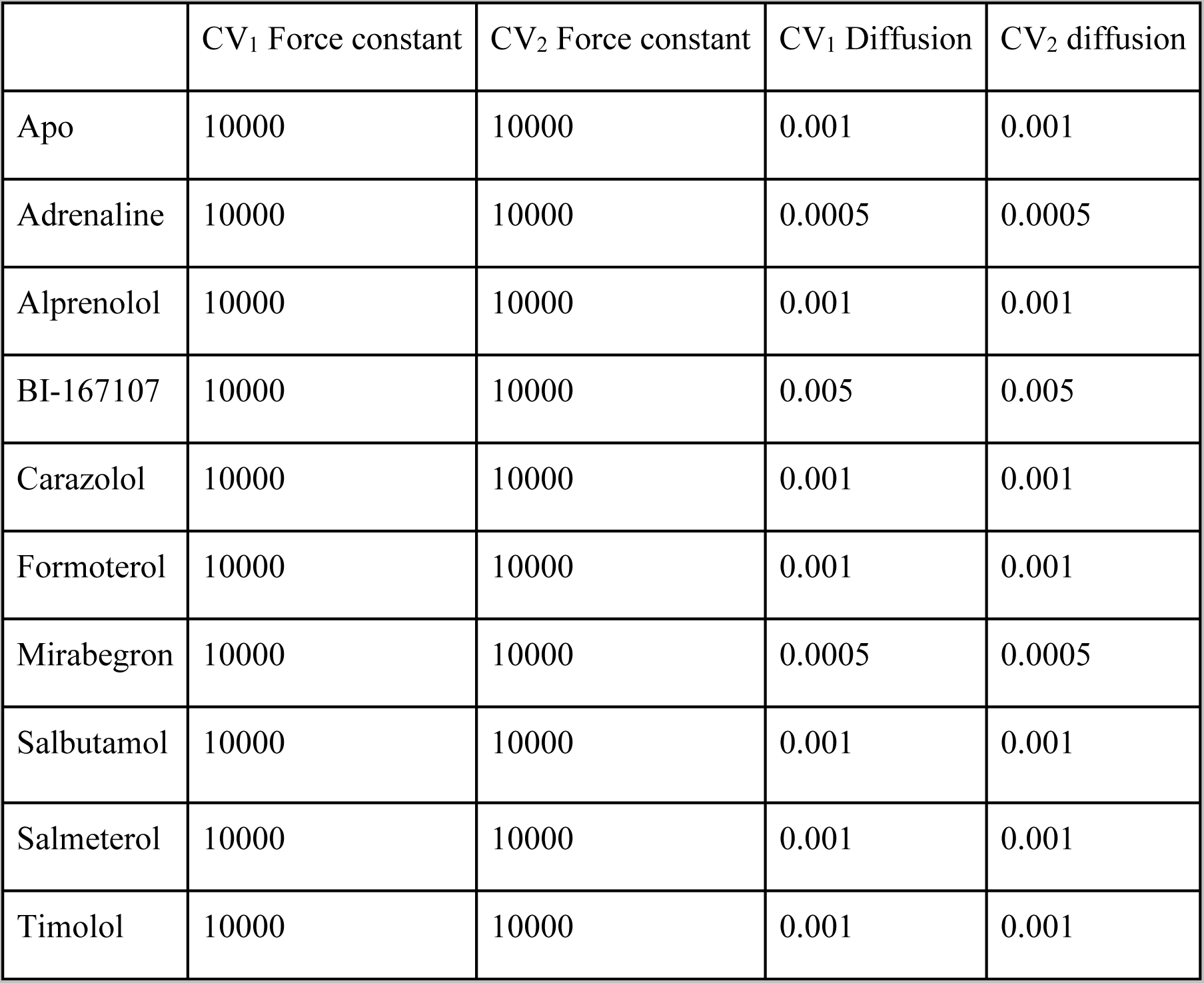
AWH parameters used for all simulations.

|  | CV<sub>1</sub> Force constant | CV<sub>2</sub> Force constant | CV<sub>1</sub> Diffusion | CV<sub>2</sub> diffusion |
| --- | --- | --- | --- | --- |
| Apo | 10000 | 10000 | 0.001 | 0.001 |
| Adrenaline | 10000 | 10000 | 0.0005 | 0.0005 |
| Alprenolol | 10000 | 10000 | 0.001 | 0.001 |
| BI-167107 | 10000 | 10000 | 0.005 | 0.005 |
| Carazolol | 10000 | 10000 | 0.001 | 0.001 |
| Formoterol | 10000 | 10000 | 0.001 | 0.001 |
| Mirabegron | 10000 | 10000 | 0.0005 | 0.0005 |
| Salbutamol | 10000 | 10000 | 0.001 | 0.001 |
| Salmeterol | 10000 | 10000 | 0.001 | 0.001 |
| Timolol | 10000 | 10000 | 0.001 | 0.001 |

The CV spaces were all defined in the range of 0.4-1.3 in CV_1_, and 0.3-1.0 in CV_2_.

### Network analysis

Collective variables need to reduce the dimensionality of the measured system for interpretability and in this case computational efficiency in converging a free energy landscape in the same space. This brings with it the risk that important degrees of freedom for moving between states may not be included in the CV description. Thus, after convergence, we prolonged the simulations (with a nearly static bias) another 100ns to achieve sufficient sampling for degrees of freedom not directly incorporated in the CV. How to exactly determine what “sufficient” sampling means is difficult to estimate even a posteriori, but we estimated that we would probably need as many coverings as would be required for convergence (∼20-50). Based on this sampling, we estimated the frame-wise free energy as a function of the collective variable (equation 2).

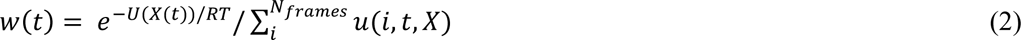

where 𝑈(𝑋(𝑡)) is the point-wise free energy estimate, 𝑅𝑇 the thermodynamic constant at 298K, and 𝑢(𝑖, 𝑡, 𝑋) reflects the binning procedure which, when summed over, gives the number of frames in the bin that is visited at time 𝑡. Moreover, 𝑋(𝑡) represents the value of the reaction coordinate at time 𝑡. Given a new reaction coordinate 𝑌, we could calculate the bin-wise energies according to equation 3.

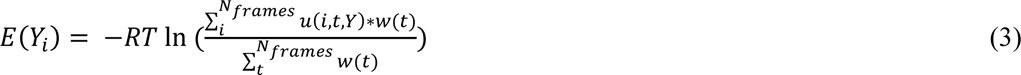

While any observable Y_i_ could be used in this formulation, we projected the free energy surface onto every minimum distance between residue pairs in the protein-ligand system. We then extracted the difference between the lowest free energy basin and highest free energy barrier to assess if there was tight or loose coupling between any alternative conformational state of each residue pair. We then used this measure as a measure of energetical coupling, which we used to construct a network via an adjacency matrix. We identified a substrate-coordinating residue on the extracellular side of the receptor, F193 as a source, and determined the most efficient coupling pathways to three sinks (P330, R131 and E268) on the intracellular side using Dijkstra’s algorithm. To get comparable estimates of a residue’s importance in this network, we calculated the betweenness centrality, the number of shortest paths passing through each residue.

### Quantification of populations

To quantify the fraction of active populations, the populations of each basin was estimated using thermodynamic weights estimated for each bin of the free energy surfaces. The basins were defined as centered in the minima and continuing up to the border of inflection points, as determined by the InfleCS methodology. Additionally, to reduce the noise from microstates, basins comprised of less than 3 bins were discarded. Moreover, the functional state of the basins (resting or active) was assigned through the following procedure: (1) The frames within each basin but after convergence was reached were extracted. (2) The snapshots were aligned with a reference active structure with PDB ID 4LDE. (3) The RMSD between the TM6 helix of each frame and the active reference structure of TM6 was calculated, where the TM6 was defined as residues 269-298. The classification was such that if the average RMSD was below 3.0Å, the basin was labeled as active, and vice versa for the resting label (see Table 2)

**Table 2.** Classification of all the energy basins for each simulation condition based on the TM6 average RMSD (as defined by residues 269-298). The cutoff value for classification was 3.0Å, criteria for active states are highlighted in red.

| Condition | Basin center CV<sub>1</sub> | Basin center CV<sub>2</sub> | TM6 average RMSD [Å] |
| --- | --- | --- | --- |
| Apo | 1.020 | 0.596 | 2.81 |
|  | 0.936 | 0.829 | 2.20 |
|  | 1.052 | 0.822 | 4.60 |
| Adrenaline | 0.907 | 0.597 | 2.10 |
|  | 1.122 | 0.681 | 4.33 |
|  | 0.848 | 0.856 | 2.89 |
|  | 0.786 | 0.685 | 1.83 |
| Alprenolol | 0.825 | 0.633 | 2.30 |
|  | 0.851 | 0.79 | 3.34 |
|  | 1.022 | 0.888 | 3.45 |
| BI-167107 | 0.801 | 0.764 | 2.55 |
|  | 1.016 | 0.695 | 2.89 |
|  | 1.049 | 0.873 | 4.65 |
| Carazolol | 1.121 | 0.794 | 5.21 |
|  | 0.986 | 0.554 | 2.17 |
| Formoterol | 0.974 | 0.698 | 2.29 |
|  | 1.010 | 0.924 | 5.21 |
|  | 1.127 | 0.690 | 3.68 |
| Mirabegron | 0.842 | 0.721 | 2.32 |
|  | 0.886 | 0.477 | 2.43 |
|  | 0.995 | 0.798 | 3.95 |
| Salbutamol | 0.889 | 0.794 | 5.34 |
|  | 0.921 | 0.521 | 2.06 |
| Salmeterol | 0.884 | 0.800 | 5.86 |
|  | 0.745 | 0.552 | 2.58 |
|  | 0.924 | 0.516 | 2.30 |
| Timolol | 0.873 | 0.831 | 4.51 |
|  | 0.811 | 0.721 | 2.97 |

In addition to simple resting/active labels, we also labeled basins based on their position in the CV space, with generally the upper right being more resting and lower left more active. With the observation of multiple potentially active basins, we also introduced the A_1_ and A_2_ states, in the lower left and lower right corners of the CV space, respectively. For the calculation of the fraction of A_1_-like population in the salmeterol and salbutamol simulations, we defined the A_1_ state as defined by the borders of the A_1_ basin in the adrenaline-bound simulations.

### Error analysis

Due to the nature of AWH simulations with an adaptive bias that changes for every iteration based on prior sampling, it is difficult to apply statistical analysis to gauge the pointwise error in the free energy estimations. Our strategy for quantifying the error was to measure the deviation from a flat coordinate distribution when the bias converges. This approach is sound because the negative bias function −U(X(r)) should approximate the underlying Hamiltonian H(X(r)), producing an effective Hamiltonian that corresponds to a flat coordinate distribution (see equation 4).

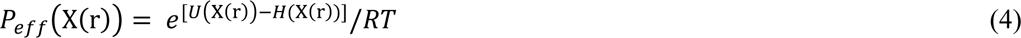

After the histogram equilibrated (meaning that the mean of the sampled distribution should be within 80% of the target distribution), we calculated the transition matrices between neighboring bins, which tell us about the transition imbalance in different regions in the free energy landscape. For an optimally converged system, all transitions should be roughly equally probable, and the higher the imbalance the higher the free energy estimate error is in this region. This can in turn be translated into free energy deviations from a flat distribution.

When basing the free energy estimation on several different walkers, it is important to check whether the sampling is consistent between them. We used the overlap metric, which is defined as the number of walkers that have a coordinate distribution of over 10% of the mean coordinate distribution. This is calculated for all points, and gives an estimation of how many walkers sample a flat distribution according to the shared bias function.

## Results and Discussion

### Computational pipeline for structure prediction and free energy landscape computation

For the methodology to be applicable, we need a starting structure (experimentally resolved or obtained thanks to structure prediction algorithms) and a deep MSA. We initiated this work from the active structure of the β2AR (PDB ID 3P0G) using an MSA of the GPCR class A family as a basis (Figure 1A) and analyzed residue pair couplings using direct coupling analysis (DCA).^21^ We then compared the coevolution coupling scores inferred from the MSA with the contacts found in the structure (Figure 1B.), where the coevolving residue pairs not in contact in the structure (false positives) are clustered according to their coupling scores using hierarchical clustering, with the number of clusters determined using the mean shift algorithm (Figure 1C.). The resulting contact clusters were then aggregated into collective variables, as weighted sums where the weights correspond to the coupling scores (and where a CV value of zero corresponds to all contacts being formed) and then used as coordinates for steered molecular dynamics to enhance the sampling along the chosen dimensions. The ensemble explored with steered MD simulations (Figure 1D) is then analyzed using a machine learning model to extract collective variables describing the conformational ensemble obtained thus far. A good CV to describe the conformational landscape should encompass the degrees of freedom with the highest intrinsic energy barriers. We thus attempt to find these degrees of freedom by finding those that are maximally separated in the maximum-margin sense between two states. Maximum-margin separation would result in a steep potential function and is thus a good approximation to finding these degrees of freedom. In this work, we used a support vector machine (SVM) with a linear kernel.^22^ The dataset is based on the tracked coevolving residue pair distances, both false and true positives, corresponding to contacts that form and break during the conformational change, respectively (Figure 1E.). This CV was then divided into positive and negative coefficients and used for AWH simulations in order to ensure a monotonic response to increasing distances in the simulations, which we have observed produces faster diffusion rates across the CV space. In principle, should the exploration not be sufficient (identified by e.g. the lack of other basins or structural features missing), the already explored degrees of freedom could be down weighted and another round of exploration could be made until sufficient conformational exploration is seen. In this work, one iteration was sufficient.

**Figure 1.**
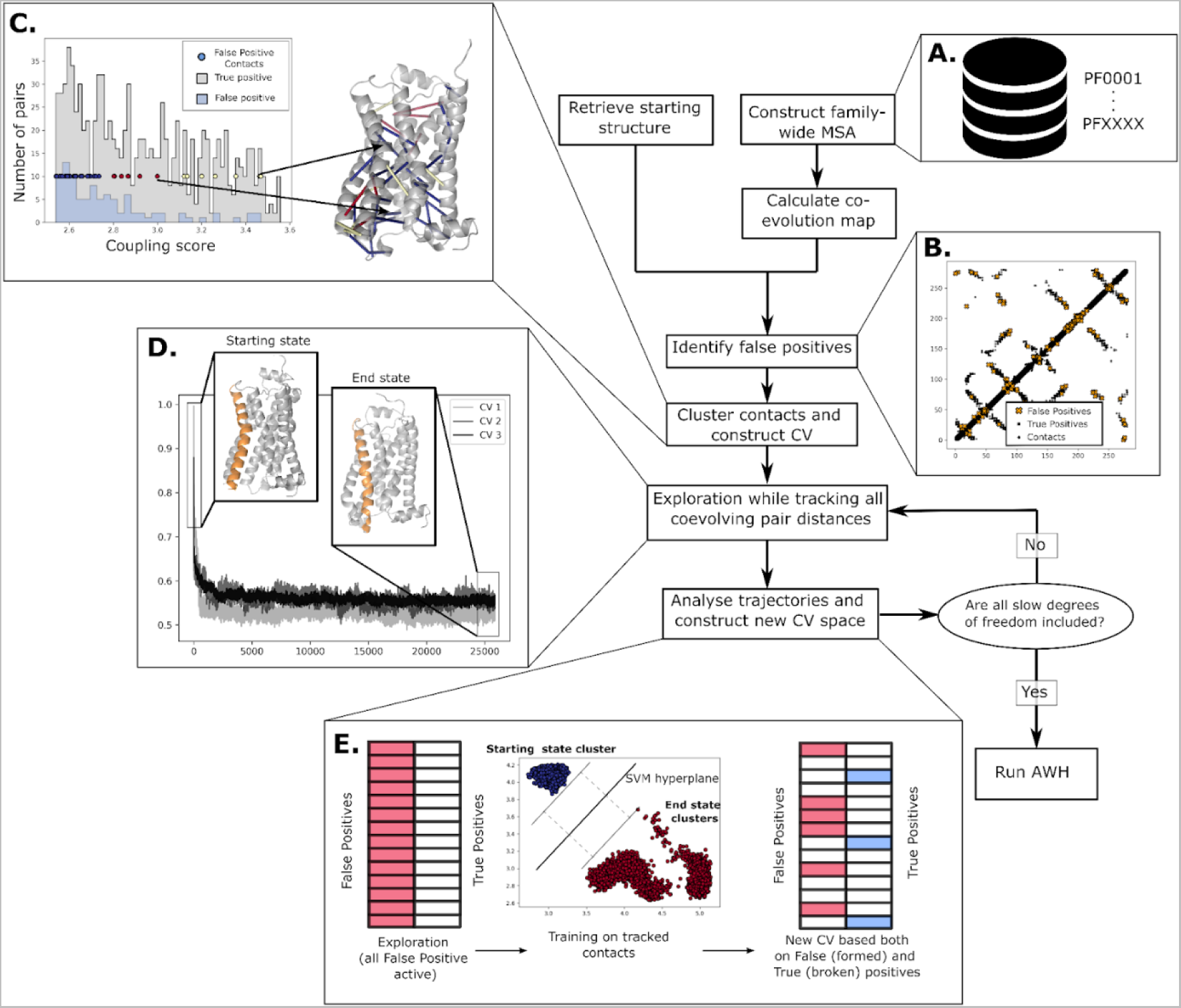
The coevolution-driven conformational exploration framework. The prerequisites include a deep MSA **(A.)** and a reference structure (PDB code used herein: 3P0G). The first step is to identify the false positive residue pairs, i.e., coevolving residues not in contact in the reference structure **(B.)**, after which these pairs are clustered according to their coevolution coupling score using the DCA algorithm as implemented in GREMLIN. Belonging to a cluster is color-coded as a circle on the coupling score plot and on the pairs shown on the molecular model. **(C.)**. The initial CVs are linear combinations of the pairs within each cluster, where the coefficients are the normalized coevolution scores. These are used in pulling simulations to achieve the required sampling **(D.)**. Machine learning methods are then applied to reduce the space of all pairs into informative pairs that formed and broke contacts during the pulling simulation **(E.).** Steps **D.** and **E.** can be iterated until all slow degrees of freedom are included, after which an AWH simulation is run to converge the conformational landscape as defined by the new CV space.

In order to probe the conformational range of the β2AR receptor, a CV space was trained as described in Figure 1. Starting from the active state based on the PDB structure 3P0G, the discovered end state after steered MD simulations (Figure 1D) features the hallmark TM6 inward movement as is expected for any GPCR deactivation.^1^ This does not, however, guarantee that the intricate movement of all the conserved microswitches is captured. To assess whether all multidimensional aspects of the inactivation mechanism were modeled, free energy surfaces were calculated in the aforementioned final CV space using the accelerated weight histogram (AWH) method^14^ (Figure 3), natively implemented in GROMACS 2022.^23^ Using the test of the flat probability distribution described in more detail in the methods section, we found that the free energy surface converged in relatively fast timescales (∼50ns, Figure S1). In most cases, a rough indication of whether the active or inactive states are favored is evident as early as 10-20 ns per walker. As discussed in our previous work on the sugar transporter family^15^, it seems that combining coevolving residues with contacts observed in structural ensembles of different states indeed effectively captures the degrees of freedom with the highest free energy barriers, and thus enables to quickly sample the conformational landscape with adaptive biasing.

### Conformational analysis

To assess and validate the accuracy of the methodology, we first aimed to characterize the accuracy of the generated structural ensemble. We did this by extracting frames from the MD trajectories corresponding to the main energetic basins. The basins that appeared in the free energy surfaces were labeled according to the TM6 position (residues 266-298) of the configurations making up the free energy minima. The latter was defined by a C-alpha RMSD metric to the reference (starting) position of the TM6 helix. Thus, we could classify the basins into inactive (above 3Å RMSD of the reference TM6 position), and active (below 3Å RMSD of the reference TM6 position) states (Table 2). Additionally, two separate basins that could both be classified as active were separately labeled as A_1_ and A_2_ based on their positions in CV space (see Methods). Since the generative procedure started with a reference structure in the active state, we assessed the accuracy of the exploration by comparing the C-alpha RMSD of the ensembles to two other reference structures (PDB ID 4LDE^24^ as an active reference, and PDB ID 2RH1^25^ as an inactive reference). The ensembles investigated were (1) equilibrium simulations of the starting structure based on 3P0G, (2) equilibrium simulations after non-equilibrium pulling and (3) the ensemble of snapshots taken from the resting basin after the AWH simulations had converged (Figure 2A.). The RMSD towards the active state is increased by the end of the exploration simulation, but only slightly decreased towards the inactive reference; however, the exploration is enough for an appropriate CV to be learned such that the correct inactive ensemble (<1.5Å RMSD) is indeed sampled during the AWH simulations. As one might expect, the non-equilibrium nature of the exploration simulations give rise to unphysical artifacts such as helices breaking or poor sidechain stacking, but this is consistently resolved on the timescales of the AWH simulations. An important result is that the basin states from the AWH simulations overlapped with the experimentally determined resting states of the receptor (Figure 2A), confirming that we can indeed find alternative states in the conformational cycle of a receptor using this method.

**Figure 2.**
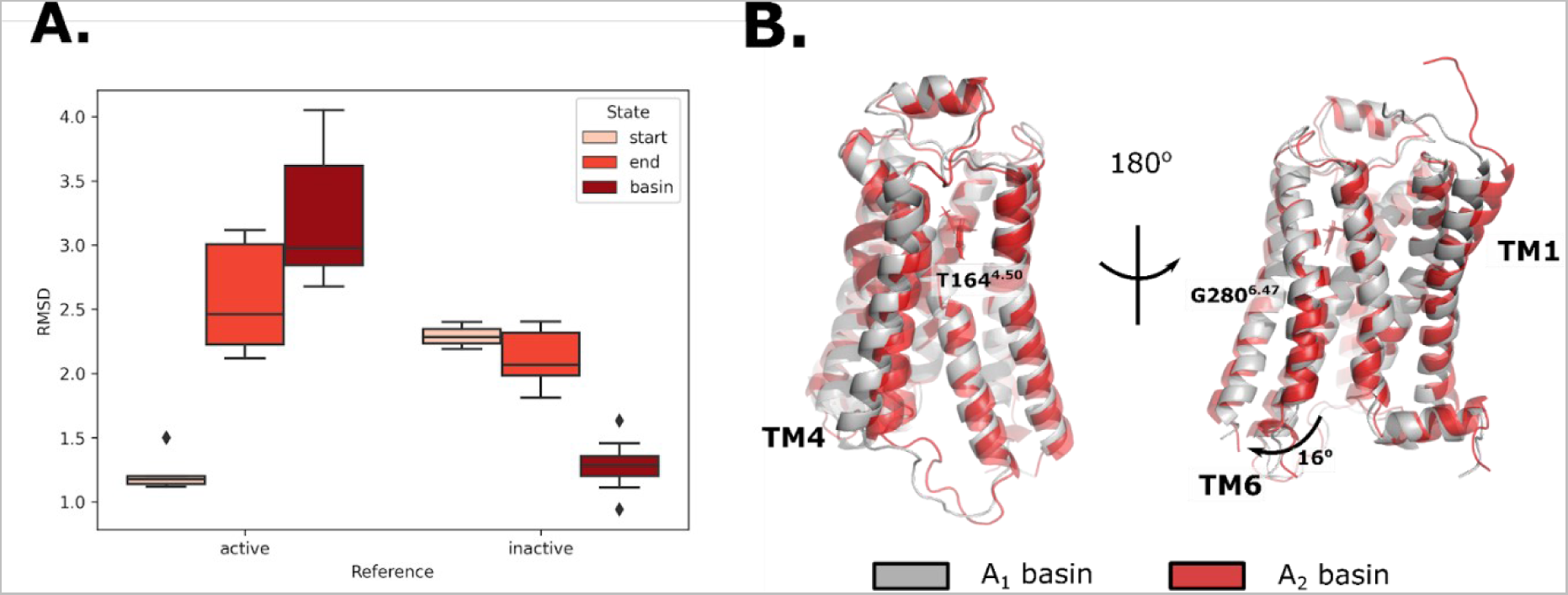
Structural analysis of the explored conformations extracted from MD simulations. **A.** RMSD towards an active (PDB code: 4LDE) and inactive (PDB code: 2RH1) reference. The structures compared to were taken from the start- and endpoints of the exploration simulations, as well as extracted from the resulting resting free energy basin from AWH. **B.** Structural differences between the active basins, divided into A1 and A2. Note that most of the differences stem from hinges in two positions in the TM6 and TM4 helices.

As the entropy of a dissociated TM6 helix in the active conformation is higher, we divided the active population into A1 and A2 states based on their position within the CV space (Figure 3 and Table 2). Upon visual inspection of the conformational ensembles in these basins, it became evident that they corresponded to alternative conformations of the TM6 helix adopting different radial positions at similar TM6-TM3 distances (Figure 2B). We found that the angle between the TM6 helix conformations was on average 16 degrees radially, and that the hinge position of this movement was located at G280^6.42^ where the TM6 helix also exhibited a slight bulge. The movement of TM6 was also observed to be accompanied with a similar hinge-driven motion in TM4 that starts with a twist at T164. Additionally, different ligands stabilized the A1 and A2 states differentially (Figure 3), which may hint at remnants of a molecular mechanism responsible for biased agonism. Interestingly, both active states are separable in this CV space even though the reference structure (4LDE) more closely resembled the A1 basin, showing that alternate local states can also be explored and sampled with this methodology.

**Figure 3.**
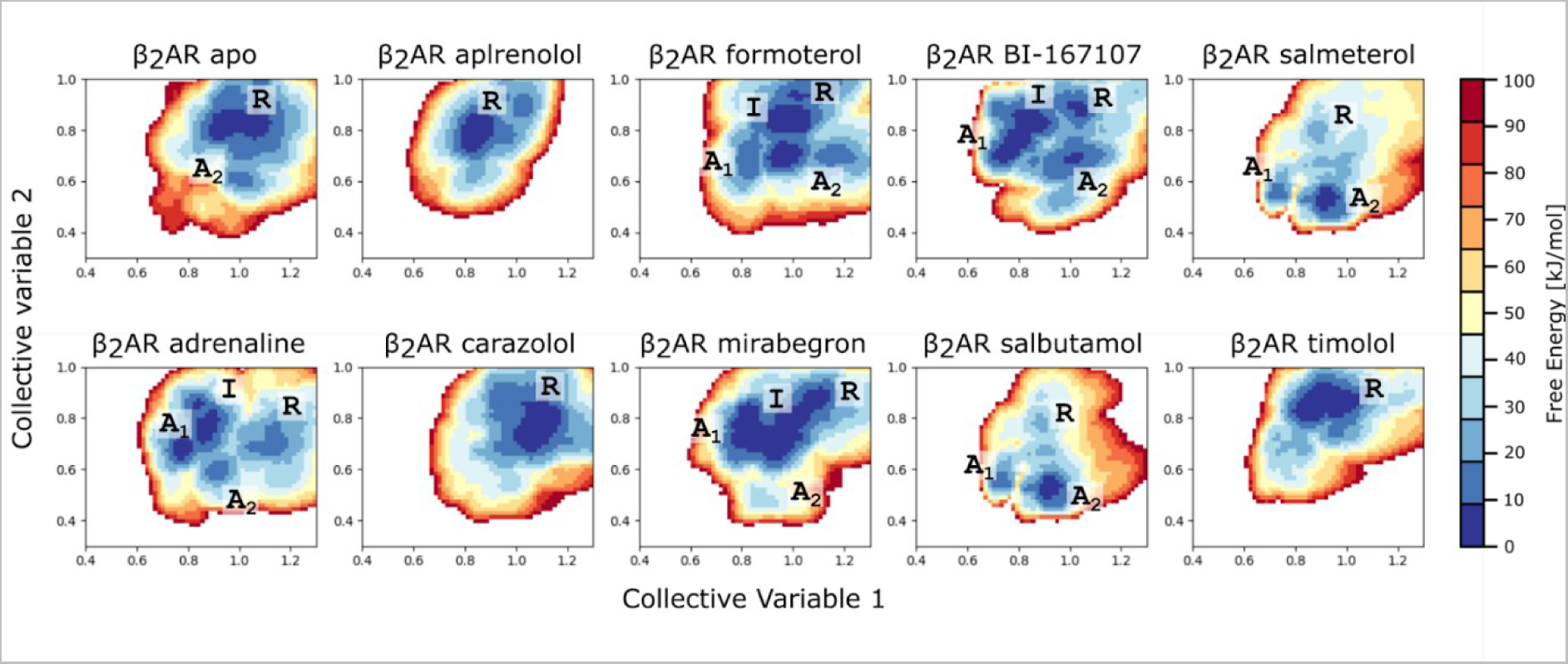
Free Energy surfaces of the conformational landscape of β2AR in the given CV space in the absence and presence of different ligands, ranging from inverse agonists to antagonists, partial agonists and full agonists. The different basins or areas of the free energy landscapes are marked with R (resting), I (intermediate), A1 (Active state 1) and A2 (Active state 2).

### Energetical analysis

Even though the correct conformational space may have been searched as indicated previously (Figure 2A), we aimed to evaluate the quality of our free energy landscapes by comparing the fraction of the β2AR population that resides in the active state. Thanks to the availability of a multitude of functional experiments measuring the efficacy of various ligands^26–32^, we could correlate our values to the %Emax values in relation to adrenaline, which was an available datapoint in all studies used.

First, we needed a global definition for assigning a functional state to each basin. Even though the CVs acted on the same atoms in each condition, the surfaces were not completely superimposable in terms of where the basins fell in CV space. Importantly, the resting and active states are separable in each of the simulation conditions even though they may be shifted. Interestingly, this suggests that there are slight differential stabilizations of molecular conformations that still correspond to similar functional states as classified by the TM6 position. This may be due to shifts in the atomic coordinates as a response to each ligand, since chemically similar ligands such as salmeterol and salbutamol produced superimposable results. This result, however, meant that any analysis of the populations had to be based on basin population, and could not be based on specific CV values. Furthermore, the basins needed to be labeled as either resting or active, after which the populations could be summed over all active states. We based this on the RMSD of the TM6 position toward the previously used reference active structure with PDB ID 4LDE (Table 2).

Visually one can clearly distinguish antagonists and inverse agonists (apo, carazolol, alprenolol and timolol) from full agonists (formoterol, adrenaline, salbutamol, salmeterol and BI-167107), where the most energetically favorable state is either resting or active, respectively. Without any further quantitative analysis, it is possible to distinguish ligands such as alprenolol and mirabegron which cover a conformational space similar to the apo simulation, and carazolol and timolol, whose basins are shifted away from any active state. Thus, it is possible to qualitatively distinguish the behavior of the receptor bound to these different ligands. Our free energy landscapes directly show the existence of multiple distinct conformations being favored or disfavored contrary to a simple two-state model. This agrees with modern observations of the existence of many functionally relevant and accessible conformations in the form of meta-stable states.^33^

However, to test the hypothesis that the CVs generated by the unsupervised conformational exploration were able to capture the intricacies of the β2AR receptor activation mechanism, we quantified the fraction of the β2AR population in all active states for each condition (Figure 4), the different activation states being defined as all bins within the inflection border of each local minimum. Additionally, we plotted these active fractions against experimentally measured relative Emax values (using adrenaline Emax levels as a reference). This ensured experiments could be compared to one another.

**Figure 4.**
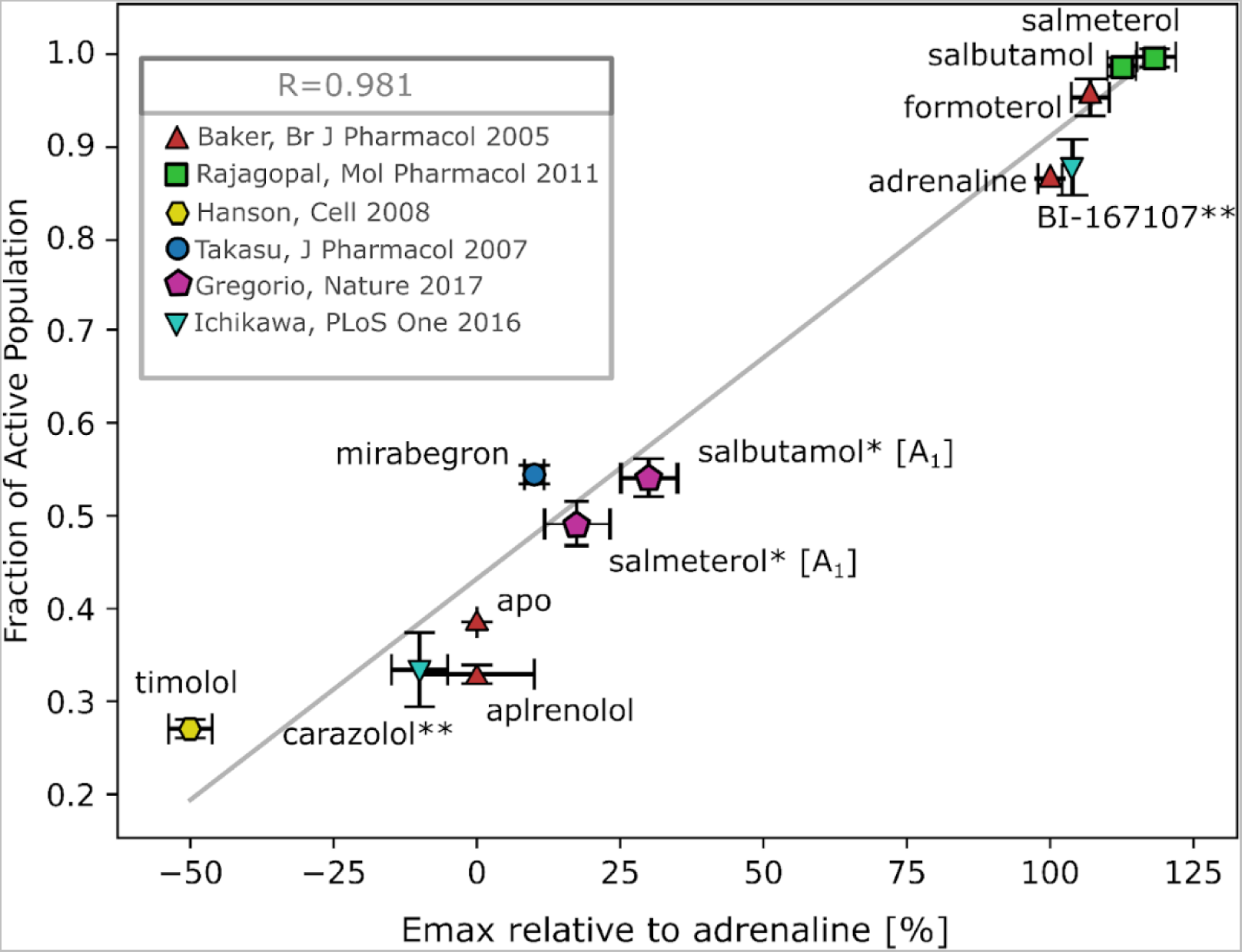
Correlation between experimental (Emax values relative to adrenaline along the x-axis)^26–32^ and computational activities (calculated fractions of the active population basins along the y-axis). The correlation line was fitted with least-squares and the R-value was calculated based on this fitted line. Due to differing efficiency values in assays measuring G-protein and β-arrestin signaling of salmeterol and salbutamol, we separately analyzed the A2- and A1 areas, which we found separately correlated with the different efficiencies. The error bars along x are taken from the references and the error bars along y are calculated based on the population difference that would arise from the calculated free energy landscape errors. * = Fraction of Active population calculated based on the population of the A1 basin, where the A_1_ basin is defined by where it is located in the adrenaline bound simulations. These points were not included in the calculation of the correlation factor R ** = Assumed computational values averaged between Ichikawa et. al. (2016)^31^ and Fleetwood et. al. (2020)^34^. These points were not included in the calculation of the correlation factor R.

Additionally, in the cases where no experimental values could be found in the literature (BI-167107 and Carazolol), we used the mean value obtained from our previous work on this receptor using string simulations.^34^

The quantification analysis shows that, in addition to displaying differentially stable resting and active states comparing full and inverse agonists, the relative populations of said basins is consistent with relative Emax values, including those determined for partial agonists (Figure 4). In contrast to previous work on similar systems, we were able to estimate the populations directly, instead of inferring dynamics from structural observables, such as the TM6 angle or microswitch positions. This suggests that the CV trained on coevolution-driven conformational exploration can indeed capture the intricacies of the microswitches that govern the ligand response from the receptor. Thanks to the correlation based on the fraction of active populations, we can conclude that ligands such as salmeterol completely activate the receptor by skewing the entire population to the active state. This suggests that the salmeterol activation of the β2AR is the theoretically highest activation response one could obtain.

In ^19^F NMR spectroscopy experiments, Kim et. al suggested the β_2_AR enthalpy changes could stem from basal activity of β2AR^35^ due to weaker interactions stabilizing the resting state than in the active state.^36^ A similar conclusion was reached using orthogonal ^19^F NMR experiments as well.^37^ The existence of basal activity is probably due to a small fraction of the receptor residing in the active state, which according to our analysis should be around 38% (Figure 4). In the same study, Kim et. al found that “two inactive states accounted for roughly 60% of the total spectral intensity for the apo and inverse agonist saturated samples”^36^, which quantifiably agrees with our estimate of 100 -38 = 62%.

Most experiments utilized the cAMP Emax value as a percentage of adrenaline activation of the G-protein pathway, while others additionally probed the Beta-arrestin pathway and quantified biased signaling.^27^ Interestingly, while most agonists stabilized active-like A_1_- and A_2_ basins, salmeterol and salbutamol exhibited the strongest shift towards one of these two basins. Curiously, these ligands have been reported as biased agonists in different experiments, where in cAMP-based assays, the activity is higher than that of adrenaline, whereas when probing the β-arrestin pathway^32^ or in FRET-based experiments^30^ they are both classified as partial agonists. Since salbutamol and salmeterol both exhibited similar and unique behavior in that the A_2_ basin was significantly more stabilized than the A_1_ basin, we wondered whether the unique behavior could be related to the biased signaling observed experimentally. We thus set out to separately quantify the relative A_1_ population for these two ligands to see if we could relate the two different basins (Figure 2B) with biased agonism. While relative population of the A_1_ basin correlates with the activation of the β-arrestin pathway (Figure 4), it should be mentioned that this is highly dependent on the definition of the functionally relevant A_1_ state.

Nevertheless, we can still qualitatively identify differentially stabilized basins, and reasonably explain biased agonism. Interestingly, this correlation leads to interesting mechanistic implications of the A_1_ and A_2_ structural snapshots of figure 2B, suggesting that biased signaling is dependent on local rearrangement of the TM4, TM1 and TM6 helices. Our observation is consistent with the model proposed by Liu et al. based on evidence for ligands favoring one of two distinct conformations.^38^ Additionally, Zhao et. al. observed that different conformers of the receptor are necessary for stable binding of Gs or Beta-arrestin, which have different binding modes.^39^ In particular, Zhao et. al. observed that ICL3 was less stabilized during β2AR-β-arrestin 1 simulations than in β2AR-Gs simulations due to a salt bridge between R239^5.77^ of β2AR and D343 of Gs. In our simulations we observed that in the A_1_ basin, the TM4 and ICL3 indeed adopts a more closed conformation than in the A_2_ basin (Figure 2B), suggesting that population dynamics of the receptor directly impairs not only signaling, but also binding of downstream pathway proteins. Our methodology is thus able to identify alternative conformational states not necessarily on the main functional pathway. However, to firmly establish the functional relevance of the A_1_ and A_2_ states, further studies on the preferential binding of G-protein and β-arrestin will be required.

### Allosteric signaling pathway analysis

Given that the CV was able to capture the activation mechanism of the β2AR, we aimed to uncover the details of the molecular mechanism, and which molecular determinants were responsible for the differential responses by different ligands. Therefore, we analyzed the structural characteristics of the entire conformational ensembles under each condition by computing inter-residue minimum distances, including interactions with any ligand along the 4x200-400ns long trajectories. Then, we projected the free energy surfaces of figure 3 onto each inter-residue distance (see methods for details). The methodology yielded the relative energetical coupling (as defined by the largest barrier between local minima along the distance) between all residue pairs in each condition. By performing a principal component analysis (PCA) on the energetical couplings for each condition, we were able to cluster the ligand-specific molecular mechanisms of activation (Figure 5A). The first two components explained 74% of the variance in the dataset and were able to cluster ligands with different effects together. Naturally, the apo simulations were unique given that no ligand was present. Nevertheless, the closest resemblance to the coupling patterns seen in the apo simulations were the inverse agonists and antagonist simulations. Two distinct clusters were found for agonist pathways, containing both full and partial agonists, suggesting two modes of signaling through the receptor.

**Figure 5.**
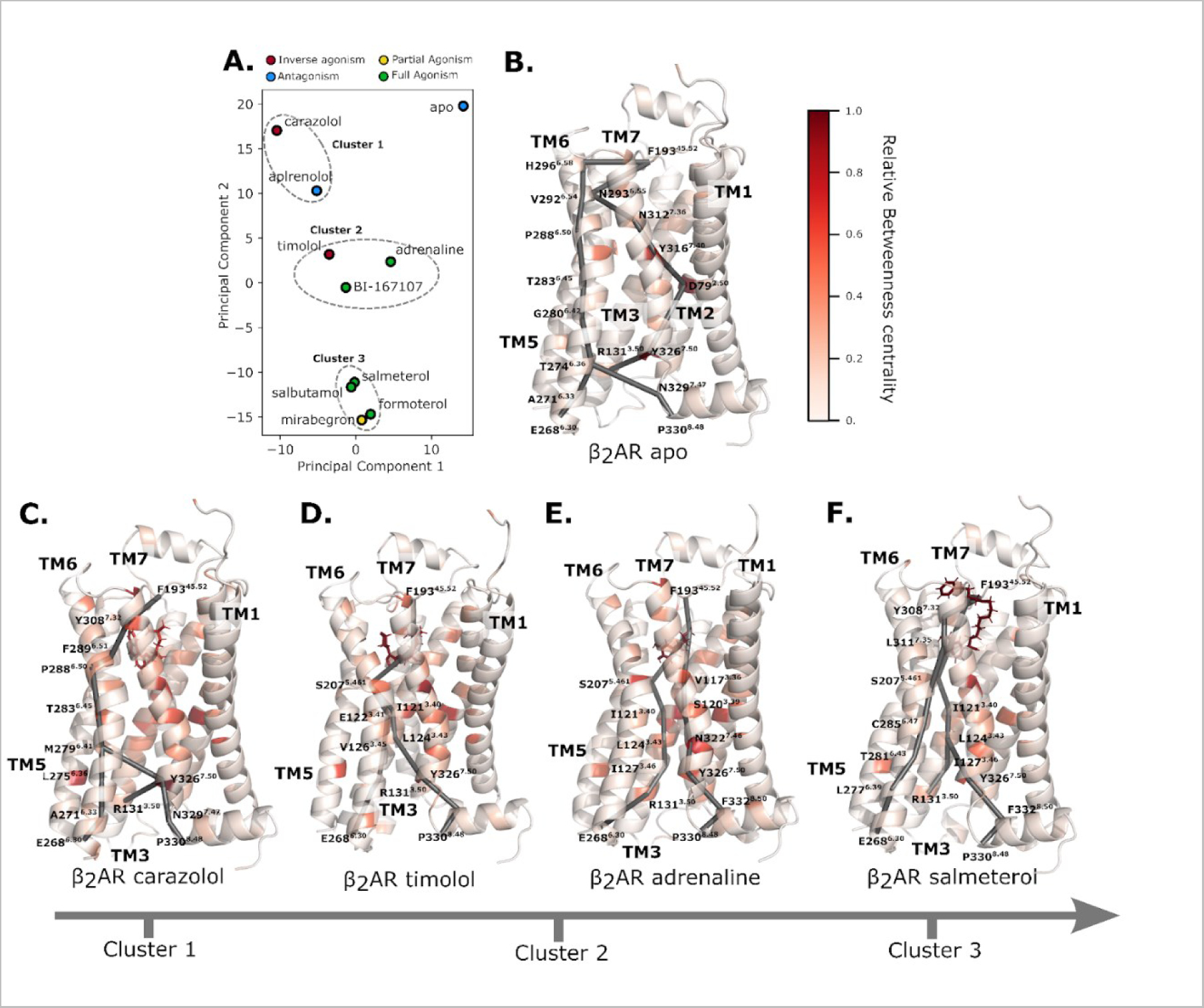
Ligands affect the activation of different allosteric networks. **A.** PCA of the per-residue energetical contributions. **B.** The β2AR apo system colored by the per-residue relative betweenness centrality, with three main allosteric paths from the ligand binding pocket (F193^45.52^) to the G-protein binding site (E268^6.30^, R131^3.50^, P330^8.48^) highlighted as sticks. Analogously plotted are the β2AR carazolol system (**C.**), the β2AR timolol system (**D.**) the β2AR adrenaline system (**E.**) and the β2AR salmeterol system (**F.**) The shortest paths shown in this figure have been made available for all ligand-bound systems in Figure S4.

In order to further analyze exactly how these coupling patterns were different, we then constructed weighted graphs representing each residue as a node and each edge as the energetical coupling scores mentioned above. We then analyzed these graphs by evaluating per-residue betweenness centrality and computing the shortest path from the extracellular to the intracellular faces of the graphs. Combining the paths and the PCA clusters, we were able to identify four main avenues of allosteric communication (Figure 5B-E). The unmodulated energetics and pathways of the receptor can be seen in the apo state.

Evidently, for a signal to be transduced from the orthosteric binding pocket (defined as F193^45.52^) to the intracellular binding pocket of the receptor (defined as three sinks at P330^8.48^, R131^3.50^ and E268^6.30^), several different pathways are exploited. While the TM6 pathway is activated in almost all cases for communication to E268^6.30^, there are alternative pathways that characteristically correspond to agonistic or antagonistic behavior. In the absence of ligand, anchoring residues around the binding site such as N293^6.55^, N312^7.38^ and Y316^7.42^ are responsible for passing the signal along to other helices in the receptor (Figure 5B). The vertical signal also passes through D79^2.50^ which has a significantly higher betweenness centrality, along with Y316^7.42^ and Y326^7.53^. Indeed, D79^2.50^ acts as a switch to bridge the signal between the two tyrosines, which sheds light on the mechanism of deactivation once the ligand has left the activated receptor. This unique mechanism of deactivation in the apo state as seen from the isolated position in PCA space (Figure 5A) suggests that the receptor activates and deactivates asymmetrically.

Unsurprisingly, the closest behavior to the apo system is found in cluster 1 containing partial inverse agonist carazolol and antagonist alprenolol. While D79^2.50^ still has a high betweenness centrality score, the communication with Y326^7.42^ occurs via TM6 as opposed to through TM2. The TM6 pathway is conserved, and the rigid communication through the backbone seems to be inherent to favoring the resting state, which also agrees with the less mobile conformation of TM6 in this state. In accordance with the lock-and-key model of receptor activation, the ligand binding residues of cluster 1 have the lowest betweenness centrality of all ligand-bound simulations. (Figure 5C) Instead, they seem to block the communication to TM7 and TM2 by not responding to the activation signal.

In both cluster 2 and 3, the signal is preferentially transduced via the ligand, indicating that the ligand has a larger energetic effect compared to the unmodulated free energy landscape. Interestingly, BI-167107 and adrenaline, two agonists, fall in the same cluster as the most extreme inverse agonist timolol (Figure 5A.). This could suggest that the energetics are communicated through similar pathways, but the energy balance is shifted to favor different states. When analyzing the details of the communication pathways in cluster 2, it becomes clear that there is a preferential signaling via TM3, in particular through a connector region centered around I121^3.40^ and L124^3.43^. Additionally, S207^5.46^ becomes a central ligand-binding site residue that transfers the signal to different pathways in all agonist-bound systems, a residue which has been indicated in the related β1AR as a determinant of agonism^40^ (Figure 5D-E). These regions with strongly state-dependent interaction patterns have been previously observed in equilibrium MD simulations by Dror et. al., where the hydrophobic connector region of I121^3.40^ and S207^5.461^ comprised two spatially distinct clusters of residues that moved together.^27^ In both the adrenaline and BI-167107 cases, the pathways split at the S207^5.461^ residue and are transduced within TM3 and TM7. Additional dynamics between N322^7.49^, S120^3.39^ and Y326^7.53^ facilitates signaling through this pathway rather than utilizing only the TM3 pathway as in the timolol-bound system. (Figure 5E) This interaction pattern corresponds to the hallmark NPxxY motif, which has been shown to be crucial microswitch, as well as functionally important and mutationally sensitive.^1, 41^

Furthermore, a clear difference between cluster 2 and cluster 1 is the absence of the TM6 pathway. In an NMR study where, among others, M279^6.41^ was probed for conformational change in the presence of an array of ligands, Kofuku et al measured a clear shift in carazolol, but no signal in the presence of agonists.^37, 42^ This is consistent with our observation that the pathway is utilized only in cluster 1 and apo simulations and passed over in other simulations. Finally, the third cluster consisting of salmeterol, salbutamol, mirabegron and formoterol exhibit more similar allosteric behavior to each other than the aforementioned clusters. In particular, there are generally two points of contact with the F193^45.52^ residue that transduce the signal through TM6 and TM3 to TM7. Additionally, Y326^7.53^, which has been shown by deep mutational scanning to be particularly sensitive to mutations,^43^ is transducing the signal via the aforementioned connector region centered around L124^3.43^ to I127^3.46^. This showcases how single residues and motifs could take on multiple roles depending on external stimuli. From an evolutionary perspective, single residues taking on multiple roles could aid multiple interactions to take place to evolve fine-tuned receptor mechanisms and provide mutational stability as a sort of safety net.

## Conclusion

Predicting alternative conformational states of membrane proteins is inherently coupled to comparing conformers that are energetically fine-tuned to exist under physiological conditions. This is further exacerbated by the fact that even a light external bias from an experimental structure determination may completely shift the delicate balance of populations, leading to complications in relying on e.g. the PDB database.^11^ In this work, we present a computational methodology to simultaneously explore and enhance the sampling of the conformational change of the β2AR receptor. By accelerating the sampling with the AWH algorithm, we also enabled on-the-fly free energy landscape estimation, providing the crucial thermodynamic validation of newly discovered states. We found excellent structural agreement between discovered conformations and available experimental structures. The fluctuations were comparable with typical equilibrium simulations of crystal structures (∼1.5Å RMSD), suggesting that we indeed reached the resting state by simulations initiated in the active state.

Upon quantifying the fraction of active population under binding of different ligands, the stunning agreement between experimental and computationally determined percentages of activation relative to adrenaline showed that we did not only describe the appropriate global rearrangements, but also achieve a detailed exploration of individual microswitch conformations. We found further evidence to support the model of a set of loosely coupled switches and an ensemble of pathways that fine-tune the behavior of the receptor, rather than a single cascade that shifts the population of the receptor.^27^

Additionally, the unsupervised nature of the conformational exploration enabled not only the discovery of alternative conformational states, but also revealed structures that plausibly relate to biased agonism identified in the salbutamol and salmeterol bound simulations. Through this observation, we could assign the functional significance of each active basin, bridging the gap between structure and function in activated ensembles. This observation stresses the importance of analyzing full ensembles of potentially undiscovered areas of the conformational space, and possibly sheds light on the discrepancy between experiments in the cases of salbutamol and salmeterol.

By combining the recent surge of computationally predicted structures in a single conformational state (using algorithms such as Alphafold2, Omegafold or trRosetta)^8–10^, with coevolution-driven conformational exploration, it would be possible to shed light on previously unvisited corners of the protein universe. Coevolving pairs are not only prevalent in alternative conformational states but may also reduce the necessary search space for protein-protein interactions or folding pathway studies using expanded ensemble MD simulations. In principle, one could repeat the exploration many times to explore and seed new states. One may use the resulting free energy landscapes to assess the energetics of the resulting conformations. Should no new states arise, one should continue the cycle. In this paper, only the CV estimation after the first iteration of conformational exploration by non-equilibrium pulling was enough for both quick and accurate convergence.

To conclude, the fast convergence times of this work (∼50-100ns per walker, i.e. 0.2-0.4µs in total simulation time per system), is around ten times faster than some previous attempts at simulating the conformational change in similar GPCR systems^44–46^, and indeed this exact system (4.3µs total simulation time per system with the string method).^34^ This study thus provides a powerful proof-of-concept of combining evolutionary data and physics through machine learning to accelerate the exploration and sampling of the conformational space to discover new conformational states. Since the methodology is not tied to this specific system or the GPCR family, this could in principle be applied to any GPCR or membrane protein family of interest. The fact that only one conformational state is needed could provide a useful way to probe the conformational landscape of proteins that are difficult to solve experimentally or are less studied in structural biology. Combining the applicability of the methodology with the fast timescales could have wide implications of using the methodology for medium-throughput drug screening or discovery.

## Acknowledgements

This work was funded by the Knut and Alice Wallenberg Foundation and the Science for Life Laboratory, the Göran Gustafsson Foundation, the Swedish e-Science Research centre (SeRC) and the Swedish Research Council (VR 2019-02433). The MD simulations were performed on resources provided by the National Academic Infrastructure for Supercomputing in Sweden (NAISS) on Dardel at the PDC Center for High Performance Computing (PDC-HPC). We acknowledge NAISS, Sweden for awarding this project access to the LUMI supercomputer, owned by the EuroHPC Joint Undertaking hosted by CSC (Finland) and the LUMI consortium. We acknowledge PRACE for awarding us access to Piz Daint, at the Swiss National Supercomputing Centre (CSCS), Switzerland.

