## Supplemental figures 1-4 for "Coevolution-driven method for efficiently simulating conformational changes in proteins reveals molecular details of ligand effects in the β2AR receptor"

### Supporting information

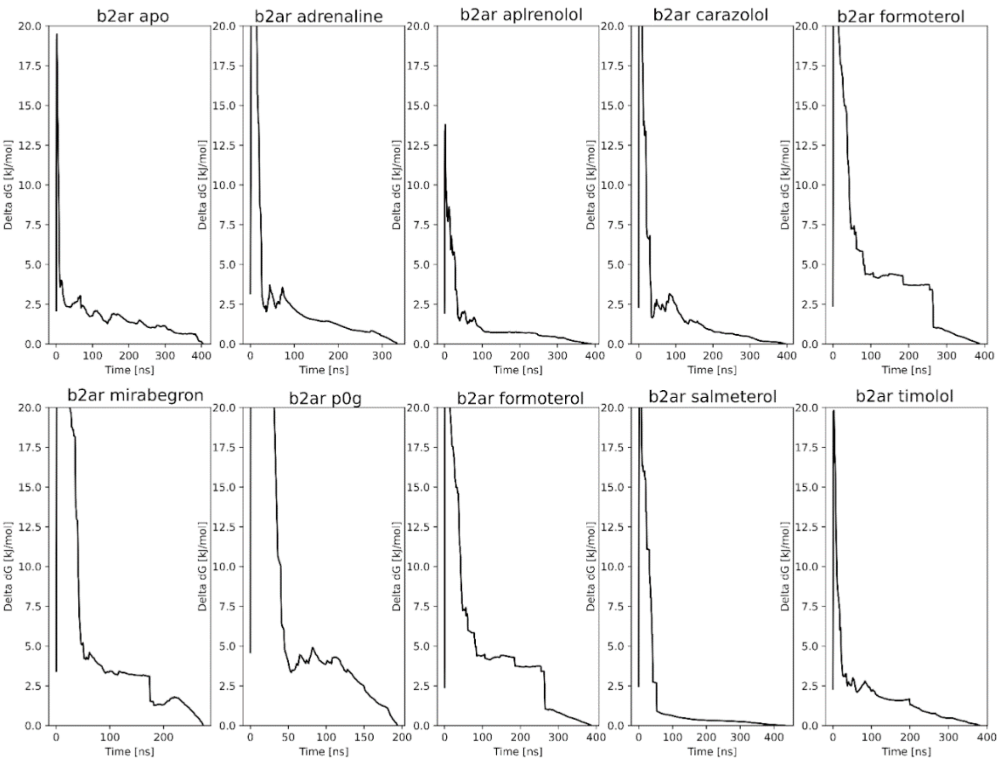


**Figure S1**. Convergence plots. Each plot shows the mean difference between neighboring frames taken over 1ns intervals over time for all simulation systems. This shows the decrease in free energy estimation in (kJ/mol)/ns over time.

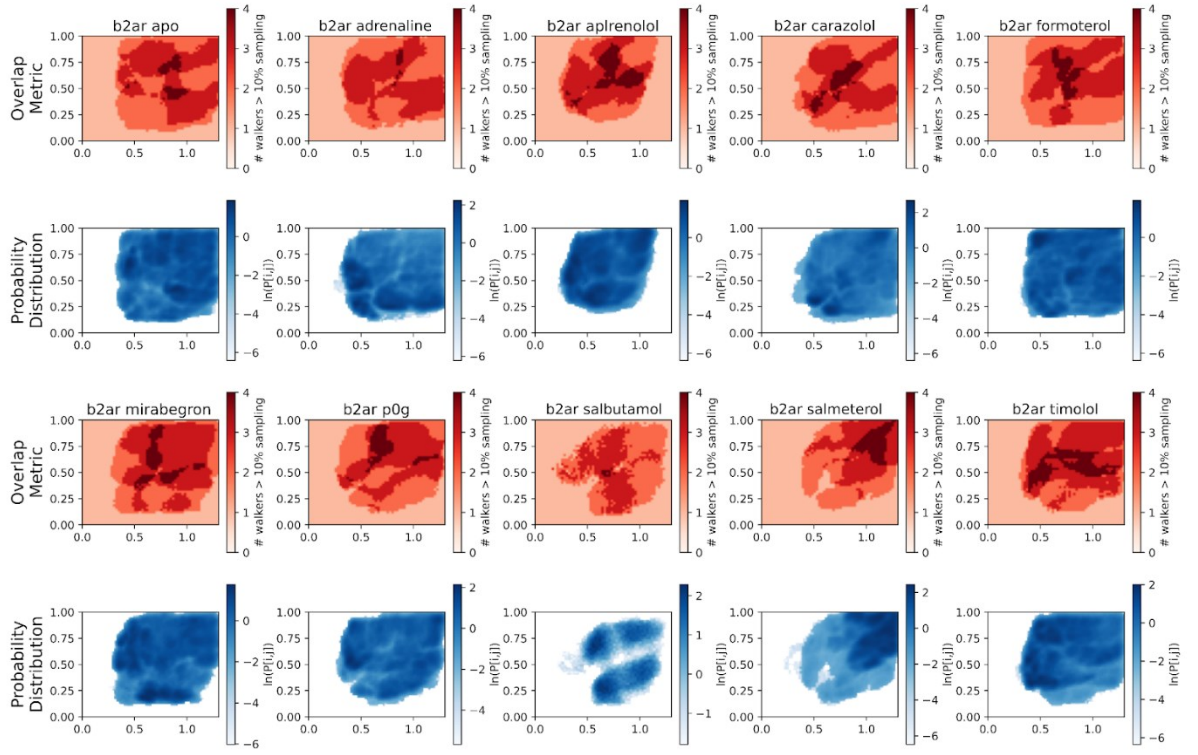


**Figure S2**. Covering plots. **Top.** The overlap metric, which shows the number of walkers (out of a total of 4 walkers) that cover each point with over 10% of the mean coordinate distribution. **Bottom.** The total probability distribution stemming from all walkers, which shows the unweighted fraction of frames spent over the entire simulation.

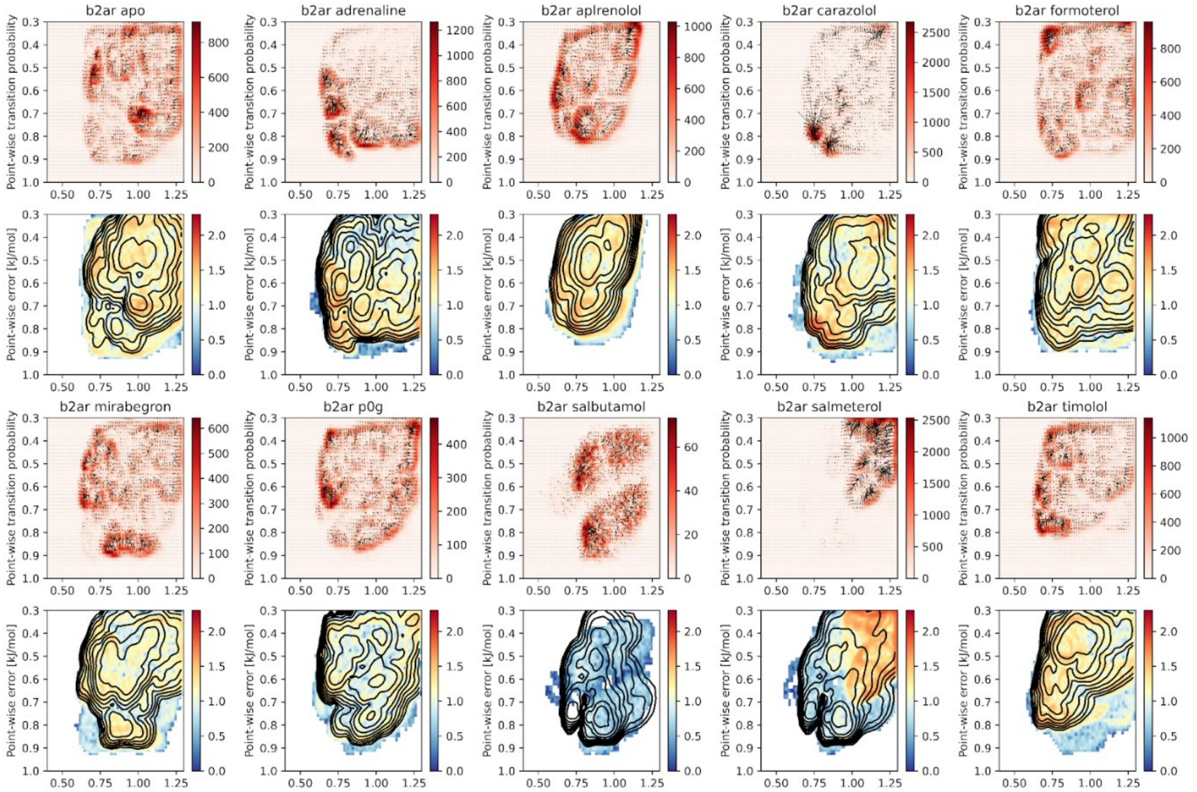


**Figure S3**. Error plots. **Top.** Vector-shaped transition imbalances colored by the magnitude of the imbalance in raw counts. **Bottom.** The estimated error, calculated as the Boltzmann inversion of the fraction of transition rates between adjacent bins. In this way, a local estimate is shown, representing the magnitude of possible deviation from a perfect bias with a flat probability distribution. The black isocurves represent the free energy surface.

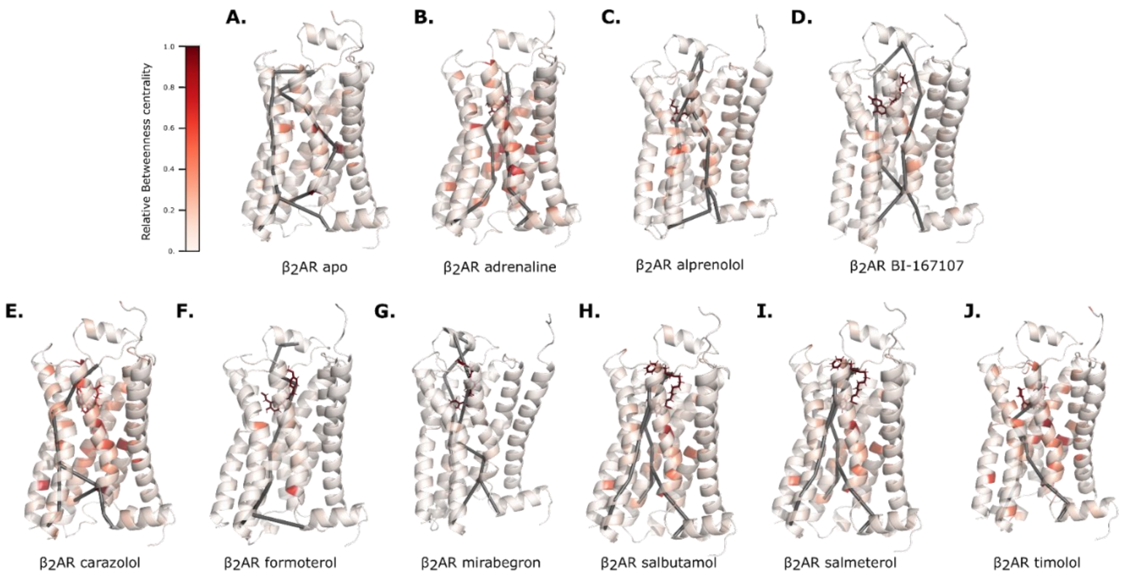


**Figure S4**. Ligand-wise network analysis. Shortest paths with three main allosteric paths from the ligand binding pocket (F193$^{45.52}$) to the G-protein binding site (E268$^{6.30}$, R131$^{3.50}$, P330$^{8.48}$) calculated with Dijkstra's algorithm and visualized in PyMOL. The networks were constructed with an adjacency matrix with undirected weights calculated as the energetical coupling according to the reweighting scheme.
